# Differences in Regional Grey Matter Volume Predict the Extent to which Openness influences Judgments of Beauty and Pleasantness of Interior Architectural Spaces

**DOI:** 10.1101/2021.03.31.437827

**Authors:** Martin Skov, Oshin Vartanian, Gorka Navarrete, Cristian Modroño, Anjan Chatterjee, Helmut Leder, José L. Gonzalez-Mora, Marcos Nadal

## Abstract

Hedonic evaluation of sensory objects varies from person to person. While this variability has been linked to differences in experience and personality traits, little is known about why stimuli lead to different evaluations in different people. We used linear mixed effect models to determine the extent to which the openness, contour, and ceiling height of interior architectural spaces influenced the beauty and pleasantness ratings of 18 participants. Then, by analyzing structural brain images acquired for the same group of participants we asked if any regional grey matter volume (rGMV) co‐varied with these differences in the extent to which openness, contour and ceiling height influence beauty and pleasantness ratings. Voxel‐based morphometry analysis revealed that the influence of openness on pleasantness ratings correlated with rGMV in the anterior prefrontal cortex (BA 10), and the influence of openness on beauty ratings correlated with rGMV in the temporal pole (BA 38) and posterior cingulate cortex (BA 31). There were no significant correlations involving contour or ceiling height. Our results suggest that regional variance in grey matter volume may play a role in the computation of hedonic valuation, and account for differences in the way people weigh certain attributes of interior architectural spaces.

## Introduction

Assigning hedonic value to sensory stimuli is a fundamental aspect of cognition (Berridge, 2018; Ellingsen et al., 2015; Skov, 2019). Hedonic values play an important role in planning and motivating behavioral responses to the sensory information (Berridge, 2004; Rangel et al., 2008; Wallis, 2007). As a general principle, organisms approach sensory objects associated with pleasure, and avoid objects associated with displeasure, pain, or disgust (Hayes, 2020; Martínez‐Garcia & Lanuza, 2018; Pessiglione & Lebreton, 2015).

Neuroimaging studies have shown that the computation of hedonic values involves neuronal populations located in the mesocorticolimbic reward circuit, including the striatum, orbitofrontal cortex (OFC), anterior cingulate gyrus (ACC), insula, and amygdala (Bartra et al., 2013; Berridge & Kringelbach, 2015; Brown et al., 2011; Sescousse et al., 2013). Sensory information serves as an important input to these processes, with hedonic valuation failing to take place if connections between perceptual and reward structures are interrupted or reduced (Loui et al., 2017; Martínez‐Molina et al., 2016, 2019).

While some categories of sensory percepts have a strong tendency to elicit either positive or negative hedonic values, no perceptual property is experienced as pleasurable or displeasurable by everyone at all times (Corradi et al., 2019, 2020). Sensory values are modulated by a long list of endogenous and exogenous factors (Coppin & Sander, 2012; Okamoto & Dan, 2013). For example, while sweet and bitter molecules generally yield positive and negative hedonic responses, including feelings of pleasure and displeasure, ingestion behavior, etc. (Sclafani, 2004; Fernstrom et al., 2012; Peng et al., 2015), how these tastes are valued vary radically with the organism’s state of satiety (Cassidy & Tong, 2017; Williams, 2014). Thus, in a state of satiety, sweet foods such as chocolate are experienced as less pleasurable, indeed sometimes unpalatable, a fact that is reflected by changes to neural activity in insula, striatum, and OFC (e.g., Kringelbach et al., 2003; Small et al., 2001). Similarly, when expectations about a stimulus are manipulated by framing information such as labels or other types of anchoring, participants report different values for identical sensory inputs (Gerger et al., 2017; Kirk et al., 2009; McClure et al., 2005; Plassmann et al., 2008). In short, hedonic values are not static or inherent to the properties of sensory objects themselves, but are rather influenced by a variety of contextual factors and individual differences (Chatterjee & Vartanian, 2014, 2016; Skov, 2019).

As a consequence of this evaluative flexibility, scientists are faced with a high degree of variability with which people judge the hedonic value of a sensory object. To make headway in understanding this variance in hedonic valuation, we recently employed linear mixed effect models to ascertain how groups of individuals differ in their hedonic response to specific sensory features (Clemente et al., 2021a, 2021b; Corradi et al., 2019, 2020). For example, Corradi and colleagues (2019) demonstrated that, while most people tested in their study preferred objects with curved contours to objects with sharp contours, variability among the participants represented 75.82% of the variance accounted for by the statistical model. Participants differed greatly in the extent to which their hedonic valuation was influenced by contour curvature: most people valued the curved contour objects more, some valued them less, and some did not take contour curvature into account in their valuation. Moreover, the extent to which participants considered contour curvature in their evaluations was consistent across time (Corradi et al., 2020) and across different kinds of objects (Corradi et al., 2019).

The issue of whether—and how—these differences in the extent to which people take account of perceptual features into hedonic valuation are related to structural or functional brain differences remains unaddressed. Identifying such relations are, however, potentially very informative about the computational processes underlying hedonic valuation, since they may help reveal mechanisms that exert a causal influence on valuation outcomes. Our goal in this study was to explore whether there is a relation between individual differences in the weighing of sensory features and individual differences in brain structure. We pursued this objective in the context of our larger ongoing research into the hedonic valuation of interior architectural spaces. Our previous studies showed that, in general, participants exhibited a majority preference for curvilinear rooms compared to rectilinear rooms (Vartanian et al., 2013), as well as for high ceiling and open rooms compared to low ceiling and enclosed rooms (Vartanian et al., 2015). These preferences correlated with differences in visual and reward‐related neural activity (Vartanian et al., 2013, 2015). However, our data also revealed that, while significant, the preference for curvilinear, high ceiling, and open rooms, was not universal (Vartanian et al., 2013, 2015). This observation raises the possibility that our group findings masked individual differences in the way these three stimulus features were represented by the individual participants and taken into account in their hedonic evaluations.

In order to test this possibility we used linear models to characterize individual differences in the extent to which contour, ceiling height, and openness influenced participants’ beauty and pleasantness ratings. We modelled *beauty* and *pleasantness* evaluations because different kinds of hedonic evaluations engage processes associated with the computation of hedonic value in different ways (Che et al., 2021). We then examined whether those measures of individual variance co‐ varied with regional grey matter volume (rGMV). We chose rGMV because it is a well‐known index of experience‐based plasticity (Kanai & Rees, 2011), and we had previously found that experience might modulate individual sensitivity to contour (Vartanian et al., 2019). To accomplish this goal, we obtained structural magnetic resonance imaging (MRI) scans from each participant, and used voxel‐ based morphometry (VBM) to calculate the correlation between variation in rGMV and individual variance in the extent to which the openness, ceiling height, and contour of interior architectural spaces determine beauty and pleasantness evaluations.

## Materials and methods

The data reported in this manuscript were collected in the same scanning session as the functional MRI data reported in Vartanian et al. (2013, 2015).

### Participants

We recruited 18 (12 women, 6 men) neurologically healthy participants (*M* = 23.39 years, *SD* = 4.49) with normal or corrected‐to‐normal vision. All participants were right‐handed, as determined by a standard questionnaire (*M* = 74.72, *SD* = 19.29; Oldfield, 1971).

### Stimuli

The stimuli for this study consisted of 200 photographs of interior architectural spaces (Figure 1). The stimuli were culled from larger architectural image databases located at the Institute for Architecture and Design, University of Aalborg, and the Royal Danish Academy of Fine Arts, Schools of Architecture, Design and Conservation School of Architecture. The stimuli were selected to represent the 8 possible combinations of openness (open vs. closed), ceiling height (high vs. low ceiling), and contour (rectilinear vs. curved contour). There were 25 images in each subcategory: 25 images of open high ceiling curved rooms, 25 images of open high ceiling rectilinear rooms, and so on. All images were standardized in terms of size and resolution.

**Figure 1.**
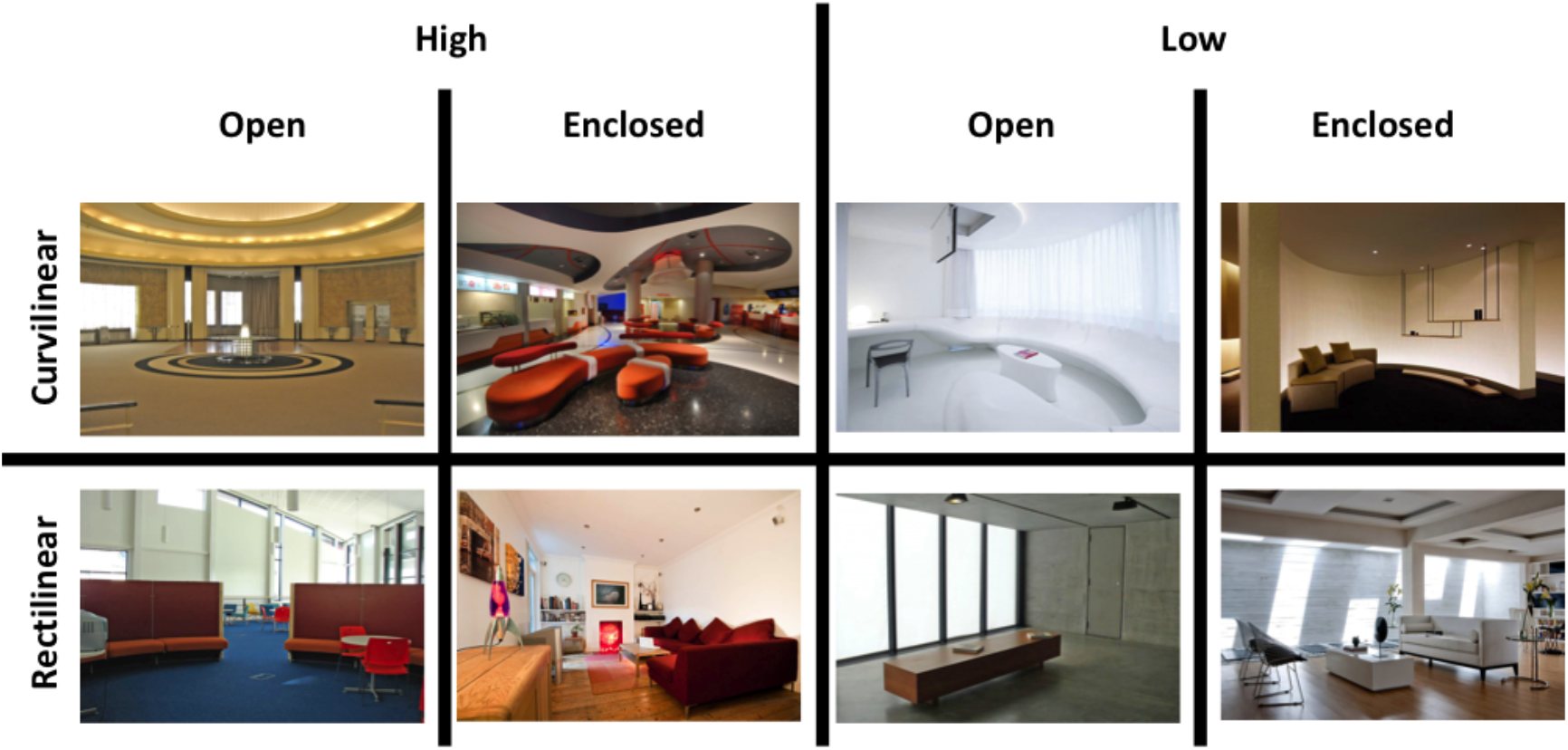
Examples of stimuli used in the study in each of the eight conditions. The figure is adapted from Vartanian et al. (2013).

### Procedures

#### fMRI acquisition

A 3‐Tesla MR scanner with an 8‐channel head coil (Signa Excite HD, 16.0 software, General Electric, Milwaukee, WI) was used to acquire T1 anatomical volume images (1.0 × 1.0 × 1.0 mm voxels).

#### Hedonic evaluation

Immediately after exiting the MRI scanner, participants were instructed to rate all the stimuli on pleasantness (using a 5‐point scale with anchors *very unpleasant* and *very pleasant*) and on beauty (using a 5‐point scale with anchors *very ugly* and *very beautiful*). The order of the ratings was randomized across the participants, as was the order of the presentation of the 200 stimuli within each block (i.e., beauty and pleasantness) for each participant. The stimuli were presented using E‐Prime software. There was no time limit for making a rating.

## Analyses

### Analysis of individual variance

Participants’ responses to stimuli in each block (i.e., beauty and pleasantness) were analyzed by means of linear mixed effects models (Hox, 2010; Snijders & Bosker, 2012). Linear mixed effects models account simultaneously for the between‐ subjects and within‐subjects effects of the independent variables (Baayen et al., 2008), unlike ANOVAs. ANOVAs usually require averaging across stimuli, which can cause the empirical Type I error rate to greatly exceed the nominal level, and lead to claims of significant effects that are unlikely to replicate with different samples (Judd et al., 2012, 2017). Linear mixed effects models provide the most accurate analyses of hierarchically structured data in which, as is the case here, responses to stimuli are dependent on, or nested within variability of individual participants. This is because they model random error at all levels of analysis simultaneously, relying on maximum likelihood procedures to estimate coefficients (Nezlek, 2001).

Linear mixed effects models are, thus, well suited to analyze preference responses, given that these often vary from one person to another and also from one object to another (Silvia, 2007). For this reason, they have been used successfully in experimental aesthetics (Brieber et al., 2014; Cattaneo et al., 2015; Mühlenbeck et al., 2015; Nadal et al., 2018; Vartanian et al., 2019). They are especially well suited to the purposes of the current study, because they provide estimates for group‐level effects, which can be compared with previous studies, and estimates for participant‐level effects, which constitute our measure of individual aesthetic sensitivity.

In the present study, two models were set up to reflect the effect of the main predictors on participants’ responses—one for the beauty ratings, and another for the pleasantness ratings. All analyses were carried out within the R environment for statistical computing, version 4.0.2. (R Core Team, 2020), using the *glmer()* functions of the ‘lme4’ package, version 1.1‐23 (Bates et al., 2017), fitted with REML estimation. The ‘lmerTest’ package, version 3.1‐2 (Kuznetsova et al., 2012), was used to estimate the *p*‐values for the *t*‐tests based on the Satterthwaite approximation for degrees of freedom, which has proven to produce acceptable Type‐I error rates (Luke, 2017). Both models included the interaction between ceiling height (*low, high*), openness (*enclosed, open*), and contour (*curvilinear, rectilinear*) as fixed effects. They also included the slope for each of these features as random effects within participants and random intercepts within stimuli. In both models, the categorical predictors were sum coded. Reference levels for the categorical variables were: *low, closed*, and *rectilinear*.

Although the models described above produce group estimates, the main aim of this study was to understand individual differences in responsiveness to visual features driving beauty or pleasantness ratings. In the linear mixed effects models, this corresponds to the modeled individual slope for each of the three features: height, openness, and contour. We thus define each participant’s aesthetic sensitivity to each of these features as the individual slope estimated from the models’ random effect structure. Therefore, after running each model, we extracted each participant’s slopes and used these values to describe aesthetic sensitivity to (a) ceiling height, (b) openness, and (c) contour, and to determine whether these individual differences in sensitivity are related to variations in brain structure as measured by rGMV.

### VBM analysis

The data were analyzed using Statistical Parametric Mapping (SPM12) (http://www.fil.ion.ucl.ac.uk/spm/software) implemented in Matlab (http://www.mathworks.com/products/matlab/). Before conducting the co‐ registration and normalization steps in SPM12, we manually reset the origin of our anatomical images (i.e., template space) using the *Display* function, with the anterior commissure as the reference point. Image registration was conducted using the *Check Reg* function and rigid‐body registration applied accordingly. All images were segmented into grey matter, white matter, cerebrospinal fluid, skull, extra‐skull, and air. Beginning with pre‐processing, the specifications for the segmentation process were as follows: we maintained at SPM12 default values for channel bias regularization (.0001), bias Full‐Width Half‐Maximum (FWHM, 60mm cutoff), and bias correction. For grey matter we maintained default values for tissue probability map and Number of Gaussians (= 1), and selected Native + DARTEL to enable image generation for DARTEL registration. DARTEL is a template creation method that increases the accuracy of inter‐subject alignment by modeling the shape of the brain using multiple parameters—in the form of three parameters per voxel. Warping and affine regularization were maintained at their default values. Smoothness (= 0) and sampling distance (= 3) were set to their default values. Field deformation was not applied. We ran DARTEL (by creating two channels for grey and white matter), following which images were spatially normalized to the Montreal Neurological Institute (MNI) brain template. This step generates smoothed, spatially normalised and Jacobian‐scaled grey matter images in MNI space. Gaussian FWHM was set to 8mm. For global normalization (the process whereby brains of different sizes and shapes are adjusted to enable group‐level inferences) we selected relative masking with a threshold of .8. We opted for the defaults of no global calculation or global normalization (the process whereby preprocessed data are scaled proportionally to the fraction of the brain volume accounted for by the represented grey matter). Subsequently, we conducted a multiple regression analysis in the GLM (General Linear Model) to capture the relation between regressors of interest and grey matter volume. Specifically, we regressed aesthetic sensitivity for each of the three features (i.e., openness, ceiling height, and contour) calculated separately based on beauty and pleasantness ratings (i.e., six regressors in total) onto grey matter volume. We also entered total intracranial brain volume calculated within SPM12 as a covariate into the analysis. Each contrast produced a statistical parametric map consisting of voxels where the *z*‐statistic was significant at *p* < .001. Reported results survived voxel‐level intensity threshold of *p* < .001 (uncorrected for multiple comparisons) and a cluster‐level intensity threshold of *p* < .05 (corrected for multiple comparisons using the False Discovery Rate [FDR]).

## Results

### Individual variance: Beauty

The results of the linear mixed effect model for the beauty ratings demonstrated that taken together, there were no significant differences between beauty ratings of low (*m* = 2.81 [2.58, 3.05]) and high ceiling rooms (*m* = 2.93 [2.69, 3.17]), β = 0.12, *t*_(170.90)_ = 0.94, *p* = .35, nor between the beauty ratings of rectilinear (*m* = 2.82 [2.59, 3.04]) and curvilinear rooms (*m* = 2.93 [2.67, 3.18]), β = 0.11, *t*_(174)_ = 0.88, *p* = .38. There were, however, significant differences between beauty ratings of open and enclosed rooms: Participants rated open rooms as more beautiful (*m* = 3.08 [2.83, 3.32]) than enclosed rooms (*m* = 2.67 [2.44, 2.90]), β = 0.41, *t*_(180.66)_ = 3.26, *p* = .001 (Figure 2).

**Figure 2.**
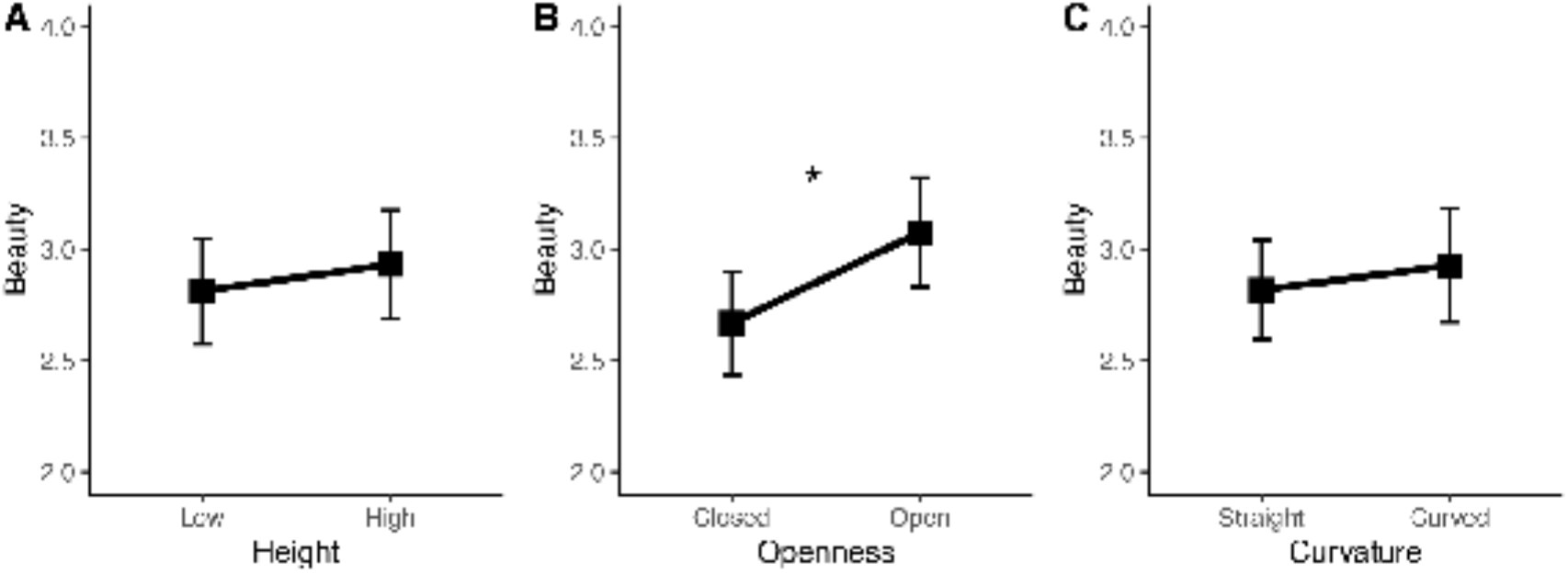
Main effects of ceiling height (A), openness (B), and contour (C), on beauty ratings. * *p* = .001.

The estimated slopes for the effects of height on each participant’s beauty ratings ranged from –0.08 to 0.47, with a mean of 0.12 and a standard deviation of 0.16. The estimated slopes for the effects of openness on each participant’s beauty ratings ranged from 0.26 to 0.79, with a mean of 0.41 and a standard deviation of 0.14. The estimated slopes for the effects of curvature on each participant’s beauty ratings ranged from –0.17 to 0.41, with a mean of 0.11 and a standard deviation of 0.15 (Figure 3).

**Figure 3.**
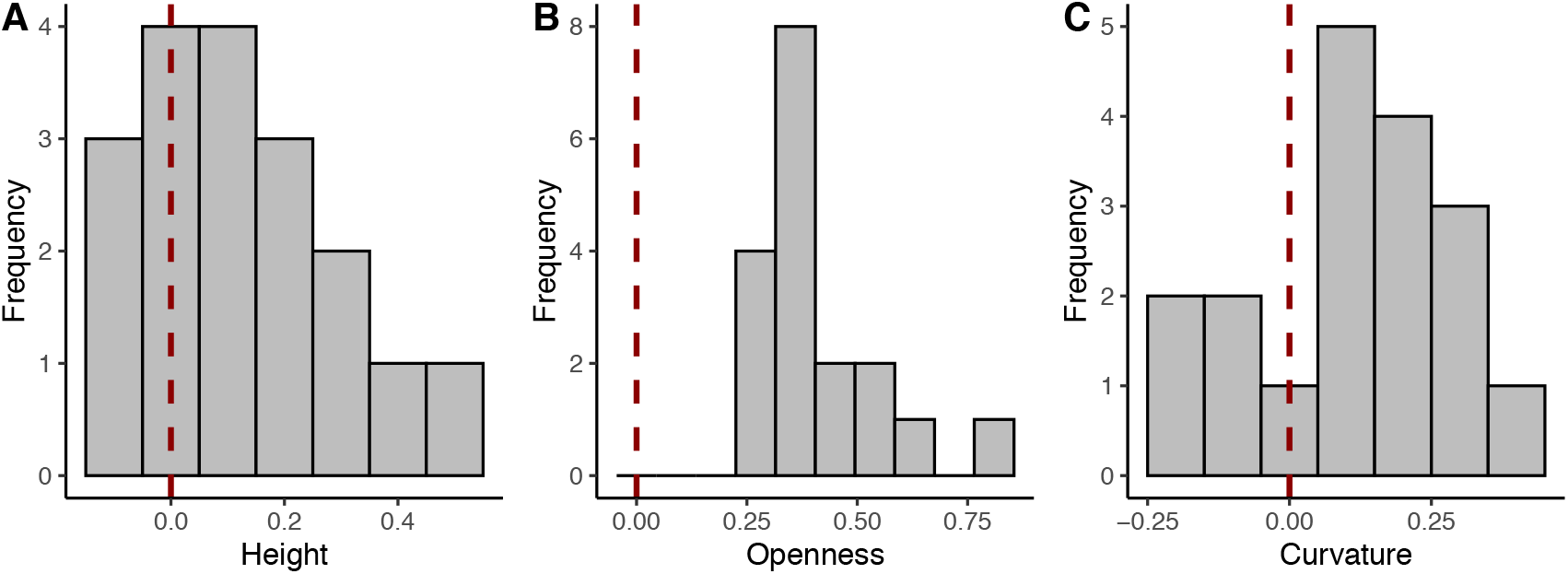
Histograms of individual slopes of beauty ratings for ceiling height (A), openness (B), and contour (C). Vertical dashed lines correspond to a slope of 0, meaning absolute indifference towards each feature. Positive slopes indicate higher beauty ratings for high ceiling, open, and curvilinear rooms. Negative slopes indicate higher beauty ratings for low ceiling, enclosed and rectilinear rooms.

**Figure 4.**
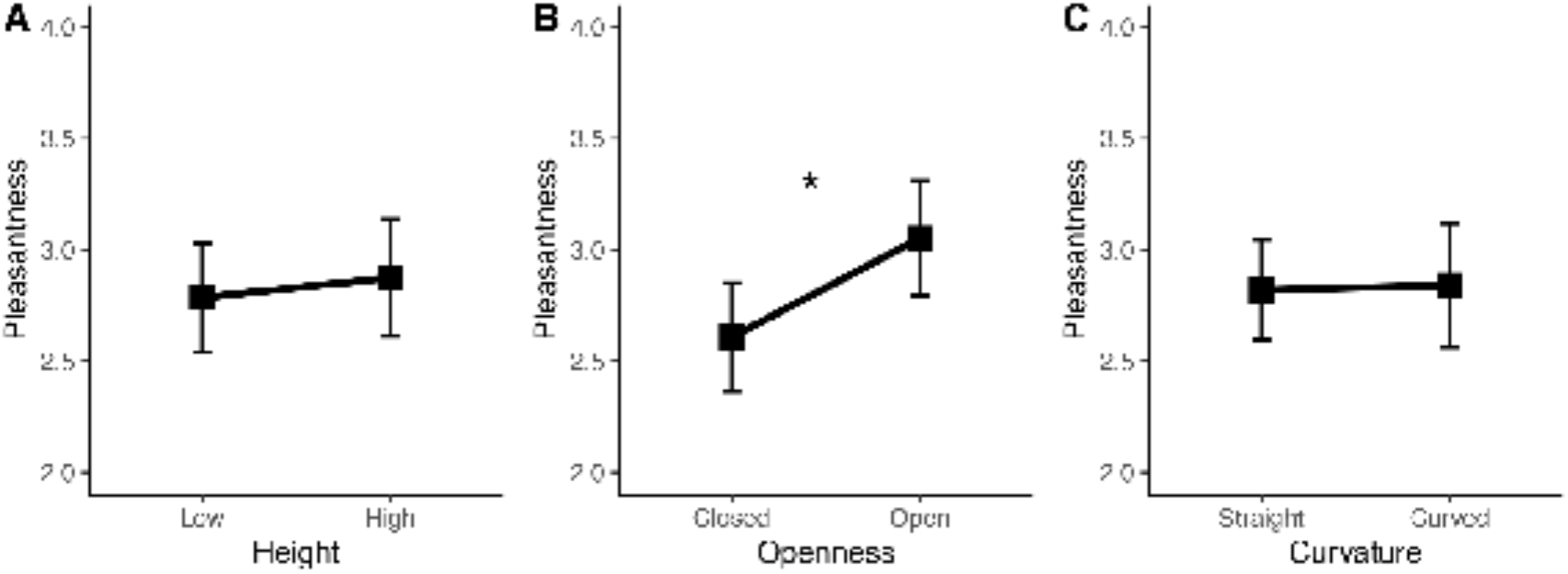
Main effects of ceiling height (A), openness (B), and contour (C), on participants’ pleasantness ratings. * *p* < .001.

**Figure 5.**
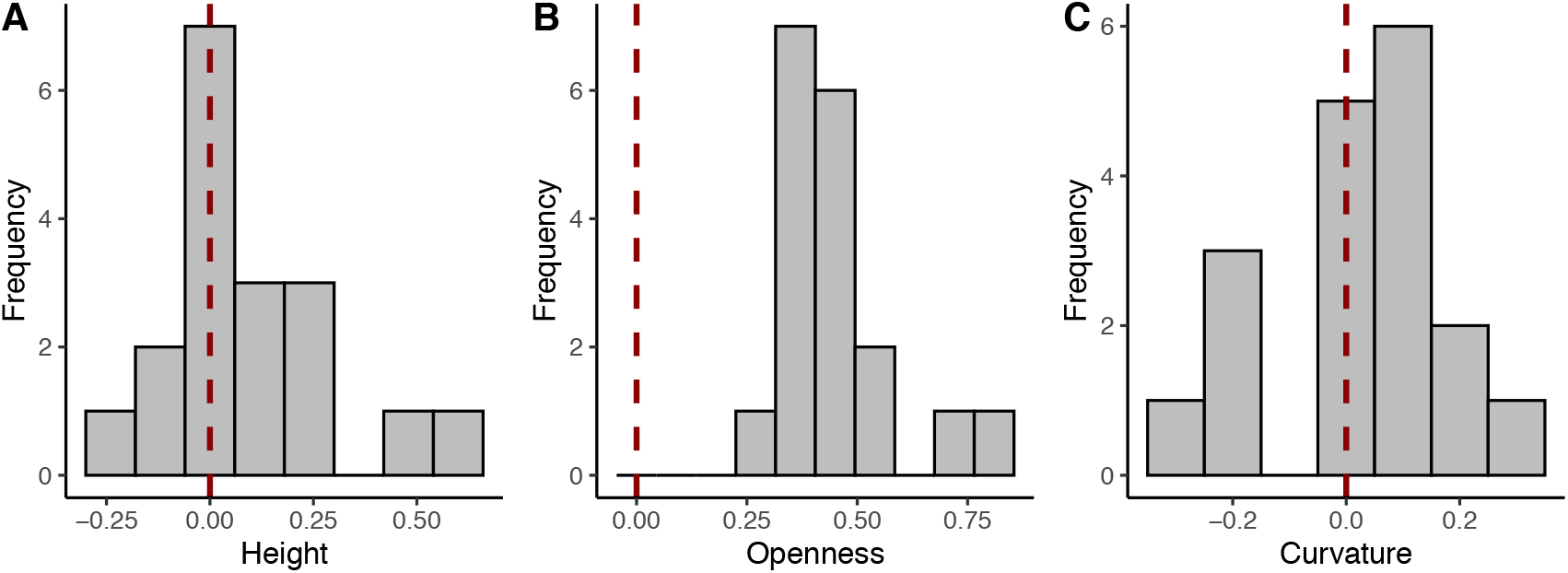
Histograms of individual slopes of pleasantness ratings for ceiling height (A), openness (B), and contour (C). Vertical dashed lines correspond to a slope of 0, meaning absolute indifference towards each feature. Positive slopes indicate higher pleasantness ratings for high ceiling, open, and curvilinear rooms. Negative slopes indicate higher pleasantness ratings for low ceiling, enclosed and rectilinear rooms.

### Individual variance: Pleasantness

The results of the linear mixed effect model for the pleasantness ratings demonstrated that taken together, there were no significant differences between pleasantness ratings of low (*m* = 2.79 [2.54, 3.03]) and high ceiling rooms (*m* = 2.87 [2.61, 3.14]), β = 0.09, *t*_(148.22)_ = 0.69, *p* = .49, nor between the pleasantness ratings of rectilinear (*m* = 2.82 [2.60, 3.04]) and curvilinear rooms (*m* = 2.84 [2.56, 3.12]), β = 0.02, *t*_(167.33)_ = 0.17, *p* = .87. There were, however, significant differences between open and enclosed rooms: Participants rated open rooms as more pleasant (*m* = 3.05 [2.79, 3.31]) than enclosed rooms (*m* = 2.61 [2.36, 2.85]), β = 0.44, *t*_(178.77)_ = 3.62, *p* < .001.

The estimated slopes for the effects of height on each participant’s pleasantness ratings ranged from –0.18 to 0.57, with a mean of 0.09 and a standard deviation of 0.20. The estimated slopes for the effects of openness on each participant’s pleasantness ratings ranged from 0.27 to 0.83, with a mean of 0.44 and a standard deviation of 0.13. The estimated slopes for the effects of curvature on each participant’s pleasantness ratings ranged from –0.31 to 0.26, with a mean of 0.02 and a standard deviation of 0.17.

### VBM results

The results from the multiple regression analysis revealed that rGMV in the right temporal pole (BA 38, *T* = 8.22, *x* = 35, *y* = 20, *z* = −36, *k*_*E*_ = 226) and left posterior cingulate cortex (PCC, BA 31, *T* = 6.70, *x* = −2, *y* = −21, *z* = 50, *k*_*E*_ = 536) covaried with individual variance in the extent to which openness influenced evaluations of beauty (Figure 6): participants whose evaluations of beauty are most influenced by openness have greater rGMV in these regions. In contrast, there was no correlation between rGMV and individual variance in the extent to which ceiling height or contour influenced evaluations of beauty.

**Figure 6.**
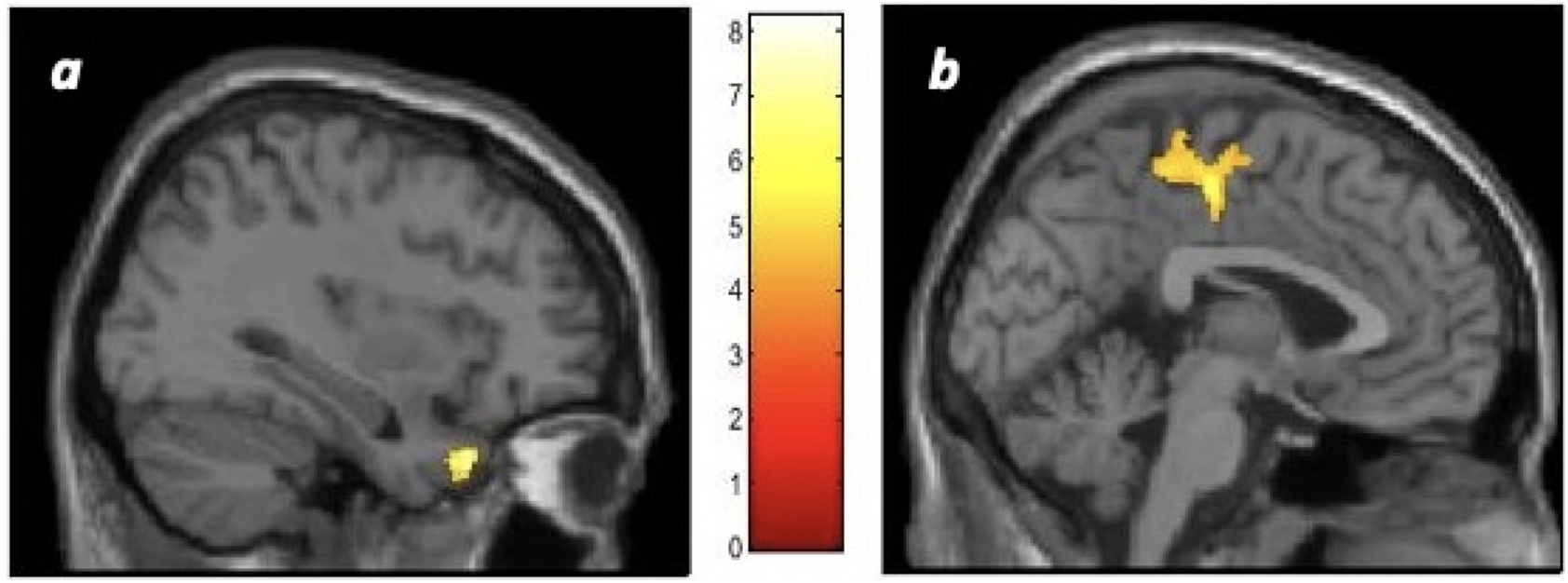
rGMV in (a) right temporal pole (BA 38) and (b) left posterior cingulate cortex (BA 31) covaried with individual variance in the extent to which the openness of interior architectural spaces influenced evaluations of beauty. The regions are overlaid on single‐subject T1 images in SPM12 and reflect the sagittal view. The bar represents the strength of the *T*‐score. The *T*‐score reflects a VBM threshold of *p* < .05 that survived a cluster‐level False Discovery Rate (FDR) correction for multiple comparisons.

In turn, the results from the multiple regression analysis revealed that rGMV in the left (BA 10, *T* = 6.78, *x* = −9, *y* = 62, *z* = 14, *k*_*E*_ = 467) and right (BA 10, *T* = 6.32, *x* = 15, *y* = 63, *z* = 9, *k*_*E*_ = 434) anterior prefrontal cortex covaried with individual variance in the extent to which openness influenced evaluations of pleasantness: participants whose evaluations of pleasantness are most influenced by openness have greater rGMV in these regions (Figure 7). In contrast, there was no correlation between rGMV and individual variance in the extent to which ceiling height or contour influenced evaluations of pleasantness.

**Figure 7.**
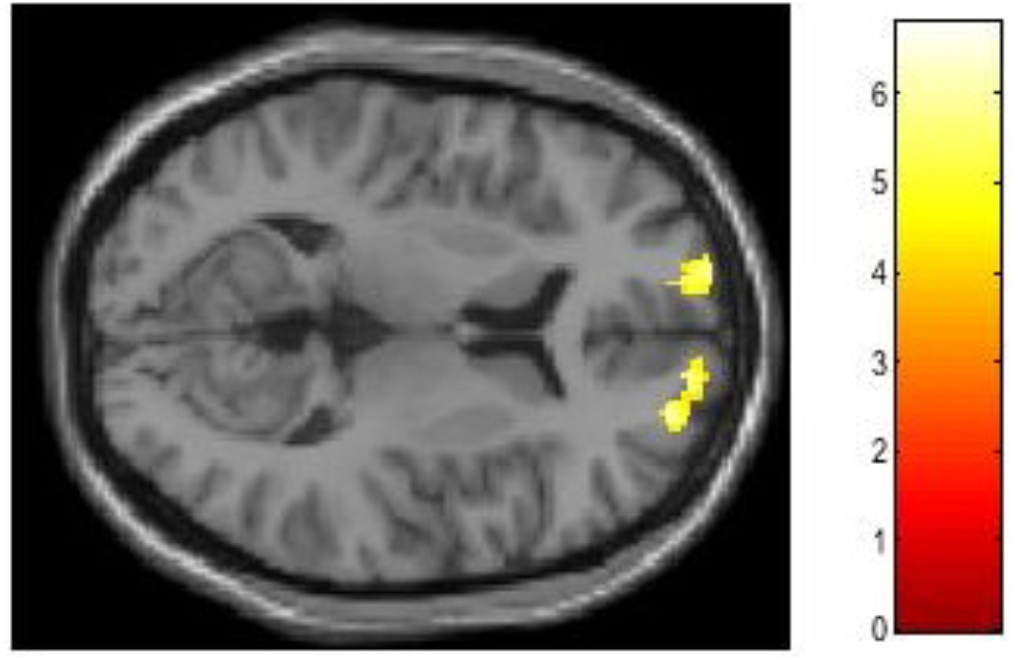
rGMV in bilateral anterior prefrontal cortex (BA 10) covaried with individual variance in the extent to which openness influenced evaluations of pleasantness. The regions are overlaid on a single‐subject T1 image in SPM12 and reflect the axial view. The bar represents the strength of the *T*‐score. The *T*‐score reflects a VBM threshold of *p* < .05 that survived a cluster‐level False Discovery Rate (FDR) correction for multiple comparisons.

## Discussion

Hedonic valuation informs value‐based decision making in a number of domains (Berridge, 2004; Hayes, 2020; Pessiglione & Lebreton, 2015). Hedonic values are modulated by contextual circumstances, including internal and external states, in order to calibrate the perceived value of a sensory object to current needs, behavioral goals, and learned expectations (Coppin & Sander, 2012; Okamoto & Dan, 2013; Ellingsen, Lekness & Kringelbach, 2015; Rangel, Camerer & Montague, 2008; Skov, 2019).

This study was conducted to test the hypothesis that individual differences in hedonic valuation can be explained as a function of variations in the way a stimulus engages mechanisms associated with hedonic evaluation. We modeled individual differences in the extent to which three visual features (i.e., contour, ceiling height, and openness) influence evaluations of beauty and pleasantness, and asked how these differences related to regional grey matter volume in the brain. We found that individual differences in the extent to which openness influences beauty evaluations correlate with rGMV in the anterior prefrontal cortex (BA 10), and individual differences in the extent to which openness influences pleasantness evaluations correlate with rGMV in the temporal pole (BA 38) and posterior cingulate cortex (BA 31). The more participants’ evaluations of beauty and pleasantness were influenced by openness, the higher their rGMV in those regions. No other correlations were found involving contour or ceiling height.

While exploratory, our results have a number of implications for our understanding of hedonic valuation and its neural correlates. First, they suggest that rGMV in the right temporal pole (BA 38) and left posterior cingulate cortex (BA 31) is related to beauty evaluations (Figure 6). Olson et al. (2007) conducted a large‐scale, systematic review of the neuroimaging and patient literatures of the temporal pole, noting that historically it has been considered by anatomists to be a paralimbic region due to its proximity to OFC and the amygdala, along with extensive connections to limbic and paralimbic regions. The authors concluded that “a general function of the [temporal pole] is to couple emotional responses to highly processed sensory stimuli. The mnemonic functions of this region allow for storage of perception–emotion linkages, forming the basis of personal semantic memory” (p. 1727). In turn, PCC (BA 31) is a core structure within the default‐mode network, and its contribution to aesthetic judgment and/or experience has been illustrated in meta‐analyses of neuroimaging studies of viewing artworks (Boccia et al., 2016; Vartanian & Skov, 2014; see also Vessel et al., 2019). In combination, the association of the temporal pole and PCC with individual differences in beauty responsiveness suggests that processes related to affect and personally‐relevant semantic and episodic memory could explain differences in the extent to which people’s beauty evaluations are influenced by the openness of interior architectural spaces.

In contrast, rGMV in the bilateral anterior prefrontal cortex (BA 10) covaried with individual differences in the extent to which openness modulated evaluations of pleasantness (Figure 7). A number of different theories have been proposed to account for the core function of this region, also known as the frontopolar cortex, which sits at the top of the prefrontal hierarchy. Christoff and Gabrieli (2000) reviewed the neuroimaging data regarding this region and concluded that the “frontopolar cortex becomes recruited when internally generated information needs to be evaluated” (p. 168). Typically, this requirement arises when a given task requires the evaluation of internally‐generated information along a relevant dimension. More recently, based on an extensive review of the human and non‐ human data, Mansouri et al. (2017) have proposed that this region’s key role is monitoring the significance of multiple goals in parallel in support of goal‐directed behaviour. Toward that end, the “frontopolar cortex does not necessarily participate in the execution of well‐learned tasks but instead collects highly processed information regarding the value (cost and benefit) of the current and alternative tasks or goals to adjust the balance between the tendency for exploitation of the current task and the drive for exploring alternative reward sources or goals in the environment” (p. 647). This function likely explains its involvement in a slew of tasks that require choosing between various alternatives that differ in value. Our results suggest that self‐relevant evaluations could be contributing factors to individual differences the extent to which people’s pleasantness evaluations are influenced by the openness of interior architectural spaces.

Together, these two findings suggest that computational nodes located in cortical structures associated with visual processing and executive control of behavior are important for representing perceptual information of relevance to the hedonic evaluation of openness. Further studies, employing interventionist techniques such as transcranial magnetic stimulation (TMS) or transcranial direct current stimulation (tDCS), could attempt to manipulate processing occurring at these nodes in order to determine the precise computational roles they play.

Interestingly, we did not find differences in hedonic valuation to correlate with grey matter variation in what are typically considered to be (early) sensory areas. This was unexpected for a number of reasons. To begin with, recent neuroimaging data have shown that low‐level visual features such as contour are processed in visual and adjacent medial/lateral surface of the temporal cortex (Yue et al., 2020). This would seem to suggest that other basic visual design features such as openness might also be instantiated in visual sensory areas. Indeed, the fusiform face area (FFA), the lateral occipital cortex (LOC), and medially adjacent regions have been shown to be activated automatically by facial beauty, even when participants are not actively asked to judge beauty (Chatterjee et al., 2009). Those data suggest that hedonic evaluation might involve computations of value in early sensory and adjacent areas, as has also been shown to be the case for architecture (Coburn et al., 2020). However, the combination of the specific construct under examination (i.e., hedonic variation) and our analytic approach (i.e., VBM) can likely explain our pattern of results. Specifically, exploration of structural neural variation in relation to various traits and capacities using VBM is likely to reveal experience‐based neuroplasticity in the brain that represents the interplay of both top‐down and bottom‐up processes (Kanai & Rees, 2011). In addition, it is possible that the extent to which specific visual features influence hedonic evaluations might be related to schemas or templates encoded in higher‐level brain regions such as the prefrontal cortex (see Wood & Grafman, 2003). According to this representational approach, even though the inputs themselves can be sensory, knowledge representations about those inputs can be stored elsewhere in the brain, as was found to be the case here. Future studies can explore differences between neural regions that *process* specific types of input (e.g., openness) using functional neuroimaging, as well as those that store their long‐term *knowledge representations* using structural brain imaging.

An important finding from the study was that the VBM results varied based on the ways in which hedonic valuation was measured, in terms of beauty vs. pleasantness. Despite the fact that empirical aesthetics is the second oldest branch in experimental psychology (Fechner, 1876), and that much work in this domain has involved the collection of various types of ratings from participants when viewing visual stimuli (e.g., beauty, liking, pleasantness, etc.), as a field we do not yet have a good handle on the engagement of the specific computational mechanisms that culminate in a given rating (Skov & Nadal, 2021). Recently, the same issue has arisen in the creativity literature where it was shown that the specific way in which a creative idea is measured (e.g., novelty vs. originality) is a reflection of the mental processes that represent it, in turn impacting its functional and structural neural correlates (Vartanian et al., 2018, 2020). In the domain of empirical aesthetics, it has been suggested that various types of evaluation (e.g., beauty vs. liking) may load differentially on cognitive vs. affective processes (Leder et al., 2005, 2012). This is not to suggest that liking and beauty are uncorrelated, but that their constituent computational components might differ. Following this logic, Che et al. (2021) have shown that generating a beauty judgment makes greater demands on cognitive resources than does generating a liking judgment, perhaps because the former necessitates a template‐matching step in the information‐processing queue that is not required for making a liking judgment. In the present study the neural correlates of aesthetic sensitivity for the same feature (i.e., openness) varied as a function of whether it was measured based on beauty or pleasantness ratings, suggesting that the lens via which one makes aesthetic evaluations might bring to bear its particular set of cognitive and affective processes.

Finally, we note that we were somewhat surprised to find that variation in rGMV only was found for variations in the hedonic evaluation of one stimulus feature—openness. Our own previous work has shown that in addition to openness, contour and ceiling height also impact beauty judgments in architecture (Vartanian, 2013, 2015). In this sense, null effects with respect to those features were unexpected. However, an important methodological difference between the present study and earlier works is that whereas in this case participants were given no time limit for completing their ratings, in our earlier work participants were given a time limit for the aesthetic judgment task. Recently, Corradi et al. (2018) examined the impact of presentation time on preference for real objects (Experiment 1) and meaningless novel patterns (Experiment 2). They found that for real objects—which is of relevance here—preference for curvilinear objects was greatest when presented rapidly, but absent when participants were given unlimited viewing time. It is possible that preference for curvilinear contour is largely driven by rapid, bottom‐up perceptual processes, the effect of which might be attenuated by top‐ down semantic effects if given sufficient time. As such, the absence of a time limit in the present study could explain why we did not observe an effect for contour on aesthetic valuation, and it is possible that an effect for ceiling height was absent for the same reason.

## Conclusions

We calculated the extent to which contour, ceiling height and openness influenced beauty and pleasantness evaluations of interior architectural spaces. Our exploratory results demonstrated that the extent to which openness influenced pleasantness evaluations was correlated with rGMV in the anterior prefrontal cortex (BA 10), and that the extent to which openness influenced beauty evaluations correlated with rGMV in the temporal pole (BA 38) and posterior cingulate cortex (BA 31). These preliminary results reinforce the importance of modelling individual variation in hedonic valuation at the level of specific design features, and reveal their structural neural correlates in the domain of architecture. Based on our findings, future interventionist approaches employing TMS and/or tDCS could further probe the precise computational roles that the regions identified here play in aesthetic evaluation of architectural interiors.

## Acknowledgments

This work was supported by the following Ministerio de Ciencia a innovación Grant TIN2011‐28146, 2011 and Ministerio Industrio, Turismo y Comercio, Avanza Grant TSI‐020100‐2010‐346 under the direction of Jose Luis Gonzalez‐Mora. In addition, it was supported in part by Servicio de Resonancia Magnética para Investigaciones Biomédicas de la Universidad de La Laguna. Portions of the findings reported here were presented at the annual convention of the American Psychological Association (APA) held in San Francisco, CA, in 2019.

